# Simultaneous association of grip effort with snack food items does not change preferences

**DOI:** 10.1101/2021.01.10.426077

**Authors:** Nadav Aridan, Tom Schonberg

## Abstract

Effort is typically considered aversive such that rewards requiring less effort are preferred over identical value rewards that require greater effort, commonly referred to as “effort discounting”. Although effort has been repeatedly shown to be weighed as a cost, there are indications to suggest that under some conditions it may increase preferences. An example for how effort affects preferences is the ‘contrast effect’ were items that follow effort production gain value. In the current study we examined whether the association of grip effort with snack food items would change preferences. In four experiments, we first identified each participant’s individual preferences of snack food items and subjective maximal grip force. We then associated different effort levels with items of similar preference. Finally, we tested for changes in preferences following the effort association. Across the four studies, effort association had no effect on preferences. Using Bayesian analyses, we conclude that simultaneous association of effort does not change preferences in our studies and call for a replication effort of previous findings.

## 1. Introduction

Physical effort is typically considered aversive such that rewards that require less effort are preferred over identical value rewards that require greater effort, a phenomena commonly referred to as “effort discounting” (Hull, 1943). Numerous studies demonstrated the cost of effort within value-based decision-making. Rodents, birds, and non-human primates have been shown to weigh effort against reward (Denk et al., 2005; Floresco et al., 2008; Phillips et al., 2007; Salamone, 1994; Stevens et al., 2005; Walton et al., 2006), and similarly do humans (Bonnelle et al., 2015; Hartmann et al., 2013; Kurniawan et al., 2010; Prévost et al., 2010; Skvortsova et al., 2014). Although effort costs have been repeatedly shown to reduce the probability of choosing the associated outcomes, several indications exist to suggest that at least in some conditions effort expenditure increased associated behavior and preferences. Festinger (1957), introduced the theory of cognitive dissonance, arguing that expectations could have an effect on the perception of effort or associated reward. For example, when an organism obtains a lower reward than expected for a particular expenditure of energy, if it is not possible to discontinue the effortful activity, there will be a tendency to retroactively attribute additional value to the activity or to its goal consequences. Several studies demonstrated this effect of effort on behavior, rats and birds that learned to associate different effort levels with visual cues or reward, spent more time in the high effort area in a subsequent effort free test and showed greater preference of the high effort associated reward (Clement et al., 2000; Johnson & Gallagher, 2011; Kacelnik & Marsh, 2002; Lewis, 1965; Lydall et al., 2010). Similar evidence exist in human studies, Alessandri and colleagues (2008), trained participants to associate discriminative shapes with the of relief of completing a hand grip. They showed that choice was shifted toward the shapes that followed higher effort. The shift in value was interpreted as resulting from the contrast between the objective value and the organism’s affective state (Zentall, 2010). However, only limited evidence exists for the change in preferences following effort. Thus, the mechanisms that determine the effect of effort learning on individual preferences remains unclear. Therefore, the aim of the current study was to examine how associating different levels of physical effort with appetitive snack food items would effect subsequent preferences.

## 2. Methods

### 2.1. Materials

The experiments were coded using Matlab (Psychtoolbox-3.0.13) and Python 2.7. A hand dynamometer (BIOPAC TSD121B) was used in order to measure grip force.

### 2.2. Participants

All participants were right-handed and had normal or corrected-to-normal vision, no history of psychiatric diagnoses, neurologic or metabolic illnesses, no history of eating disorders, had no food restrictions and were not taking any medications that could interfere with the experiment. Participants were told that the goal of the experiment is to study food preferences and were asked to refrain from eating 4 hours prior to arrival to the laboratory (Plassmann et al., 2007). Participants gave informed consent, and the study was approved by the ethics committee at Tel-Aviv University. Exclusion criteria were set as insufficient range of willingness to pay (WTP; see Experimental procedure). See Table 3.1 for experiments sample sizes and demographics.

### 2.3. Experimental procedure

See Table 2 for summary of each experiment parameters.

**Table 1.** Experiments sample sizes and demographics

| Experiment | N (excluded) | Mean age [range] | Sex (male) |
| --- | --- | --- | --- |
| 1 | 20 (0) | 24.0 [18 33] | 3 |
| 2 | 27 (1) | 24.4 [19 35] | 8 |
| 3 | 29 (5) | 27.3 [19 39] | 16 |
| 4 | 24 (0) | 24.4 [19 35] | 4 |

**Table 2.**
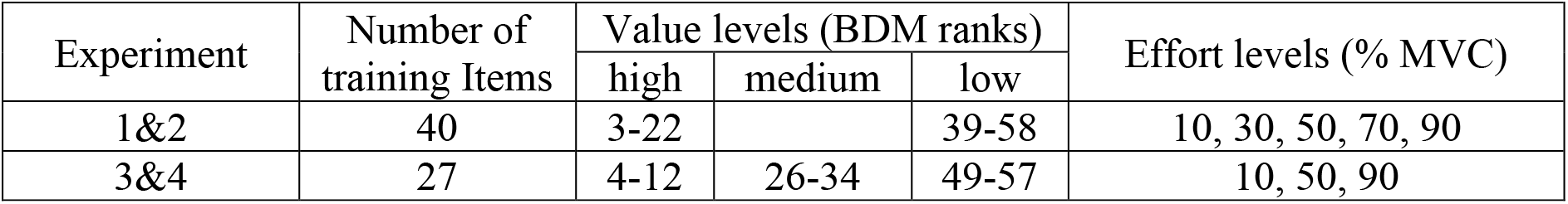
Experiments design

**Table 3.** Effects of effort and value on WTP

| Experiment | Value |  | Effort |  | Value*Effort |  |
| --- | --- | --- | --- | --- | --- | --- |
|  | F<sub>(df)</sub> | p | F<sub>(df)</sub> | p | F<sub>(df)</sub> | p |
| 1 | 17.15<sub>(1,19)</sub> | <0.001 | 1.79<sub>(4,76)</sub> | 0.14 | 1.42<sub>(4,76)</sub> | 0.23 |
| 2 | 18.25<sub>(1,25)</sub> | <0.001 | 1.88<sub>(4,100)</sub> | 0.12 | 1.25<sub>(4,100)</sub> | 0.30 |
| 3 | 8.03<sub>(2,46)</sub> | <0.001 | 1.07<sub>(2,46)</sub> | 0.35 | 1.01<sub>(4,92)</sub> | 0.40 |
| 4 | 13.99<sub>(2,46)</sub> | <0.001 | 1.42<sub>(2,46)</sub> | 0.25 | 0.18<sub>(4,92)</sub> | 0.95 |

#### 2.3.1. BDM Auction

First, participants took part in an auction in which photographs of 60 appetitive snack-food items were presented in random order. We followed the BDM procedure (Becker et al., 1964) in which participants were endowed with 10 NIS and told that they would have an opportunity to use them to buy a snack at the end of the session. During the auction, participants were presented with one item at a time on a computer screen. They placed their bid by moving the mouse cursor along an analog scale that spanned from 0 to 10 at the bottom of the screen. The auction was self-paced, and the next item was presented only after the participant placed his or her bid. This procedure has been shown to reliably provide a measure of WTP per item. An exclusion criteria was set as insufficient range in bids to properly categorize the stimuli using the method explained in the next section (over 40 bids under 1 NIS; Figure 1.A).

**Figure 1.**
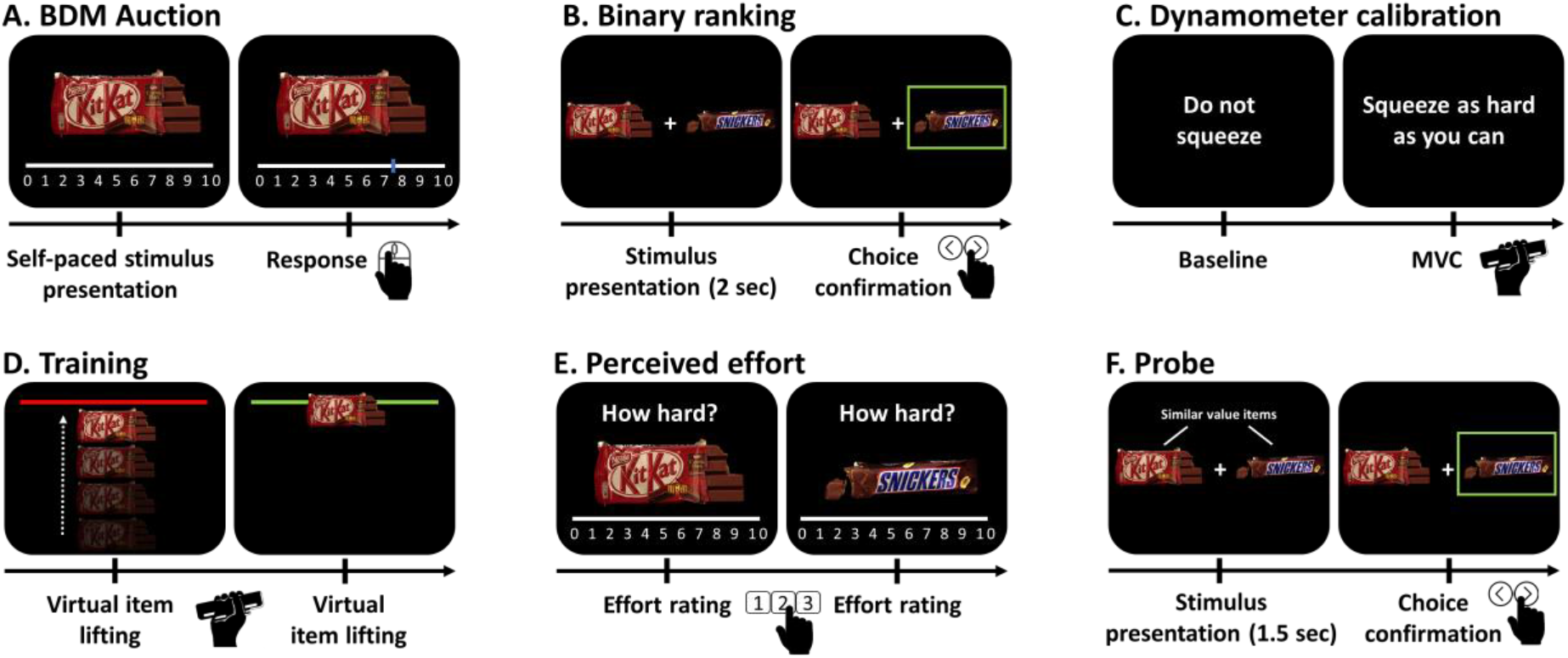
Experimental procedure. A, BDM auction. B, Binary ranking. C, Dynamometer calibration. D, training. E, Perceived-effort rating. F, Probe.

#### 2.3.2. Item selection

Items were ranked based on WTP, where the first rank is the item with the highest bid. Items that served as stimuli for the remainder of the procedure were selected based on predefined ranks that formed different value categories (experiments 1&2 N_items_=40, high value ranks: 3-22, low value ranks 39-58; experiments 3&4 N_items_=27, high value ranks: 4-12, medium value ranks: 26-34, low value ranks 49-57). Within each value category, each item was consistently associated with one effort level, keeping a similar average rank in each resulting effort category (experiments 1&2: 10, 30, 50, 70, 90 10, 50 % of MVC; experiments 3&4: 10, 50, 90 % of MVC). This selection procedure ensured pairing of similar-value items of all effort levels. These pairs made up of two items of similar WTP later presented at probe (see below) such that participants should a priori be indifferent in a choice between them based on initially stated values.

#### 2.3.3. Binary rankings (experiments 3&4)

In addition to the BDM auction, the 27 selected items, were compared with each other using binary choices to construct the initial preference ranks (each of the 27 items appeared with each of the other 26, 351 comparisons in total). On each trial, participants were presented with two items side by side and had 2 seconds to choose the one they prefer using the keyboard. Participants had 2 seconds to respond with either of two buttons on the keyboard corresponding to the left or right locations on the screen. The chosen item was highlighted with a green rectangle around it. If the participants did not choose within the allotted time, a message appeared on the screen asking them to please choose faster followed by the inter-trial fixation cross and the next trial began (Figure 1.B). Based on the assumption of choices transitivity from rational choice theory (Von Neumann & Morgenstern, 1944), pairwise binary choices were transformed into quantified ranking scores. To maximize ranking accuracy, we used the Colley Matrix algorithm (Colley, 2002), designed to solve ranking problems with relatively small number of trials.

#### 2.3.4. Dynamometer calibration

Prior to training, the required effort was calibrated to each participant’s individual maximal voluntary contraction (MVC). Participants held a handgrip dynamometer in their right hand and were instructed to produce three 2-second squeezes as hard as they could. MVC was calculated by averaging the three squeezes minus the maximal value of the baseline (Figure 1.C).

#### 2.3.5. Effort-Item association

The participants underwent a training session in which they learned to associate one effort level (experiments 1&2: 10, 30, 50, 70, 90 10, 50 % of MVC; experiments 3&4: 10, 50, 90 % of MVC) with each of the selected items. On each trial one snack image appeared at the bottom of the screen and a red bar appeared at the top. Participants had 1.5 seconds to squeeze the dynamometer as hard as needed in order to raise the snack image up to the red bar. If they managed to reach the bar, they received a feedback as the bar turned green. The amount of effort required for each item was not explained explicitly to the participants and was learned through experience (Figure 1.D).

#### 2.3.6. Perceived-effort rating

Following each item virtual lifting (experiments 1,2&3; or six training runs in experiment 4), the participants were asked to rate the item’s perceived effort. The participants were presented with a visual analog scale (VAS) and asked to rank, using numbers on the keyboards, how difficult was the current trial between 0-10 (0 for “no effort at all” and 10 for “very difficult”; in experiment 4: scale of 0-4, this was followed by another six training runs and a second perceived-effort rating; Figure 1.E).

#### 2.3.7. Fractal images rating

As a short break between the training phase and the probe, participants were presented with sixty fractal images, one at a time. They were asked to rate each one on a scale from 0 to 10 according to how much they like them.

#### 2.3.8. Binary choice probe (experiments 3&4)

Participants were presented with pairs of items. They were told that a single trial will be drawn at random at the end of the session and their choice on that trial would be honored (i.e. they would receive the item that they had chosen on that trial at the end of the experiment). We presented unique pairs of items during the probe phase. The main goal of our analysis was to test how the effort learning training affected participants’ preferences between items that had similar initial value and different associated effort level. Participants were presented with 27 unique pairs (each of the three items of a certain effort level was paired with each of the three items of the different effort level) for each value category (81 total). If the manipulation did not affect participants’ valuation of the items, they should be indifferent between them.

At trial onset, the two items in a pair were presented directly to the right and left of a fixation cross. Participants had 1.5 seconds to respond with either of two buttons on the keyboard corresponding to the left or right locations on the screen. The chosen item was highlighted with a green rectangle around it. The choice confirmation remained on the screen for 500 ms until a fixation-cross appeared during the inter-trial interval for an average of 3 s (range 1–12 s). If the participants did not choose within the allotted time, a message appeared on the screen asking them to please choose faster followed by the inter-trial fixation cross and the next trial began. Each of the 81 unique pairs was presented twice across the two probe runs. The presentation order and locations (left/right) of the items on the screen were randomized across participants and across the two runs (Figure 1.F).

#### 2.3.9. Second Binary ranking (experiments 3&4)

A second binary ranking, similar to the first one took place.

#### 2.3.10. Second auction

A second auction, similar to the first one took place. Difference in willingness to pay (ΔWTP) was calculated by subtracting the bid of the first auction from the bid of the second Auction.

#### 2.3.11. Perceived effort recognition (experiments 3&4)

Participants were presented once again with the perceived effort rating task.

### 2.4. Statistical analysis

#### 2.4.1. Null hypothesis significance testing (NHST)

All mixed linear regression analyses were implemented in R (R Core Team, 2019; version 3.6.1) with lme4 package (Bates et al., 2015).

#### 2.4.2. perceived and required effort

In each experiment, for each participant, perceived effort of each effort level was calculated as the median of the participants’ effort ratings of items from the relevant effort level. We then fitted a linear regression with perceived effort as the dependent variable, required effort as independent variable and participant as a random effect.

#### 2.4.3. Difference in WTP

In each experiment, for each participant, we calculated the difference in WTP (Δbid) for each item before and after effort association. We then averaged the Δbid across items of the same value-effort category. Difference in the mean Δbid between the different effort and value levels was tested using a two way repeated measures ANOVA.

#### 2.4.4. Binary choice probe

In experiments 3 and 4, choice ratio of higher effort over lower effort items tested using a binary logistic regression with participant as a random factor.

#### 2.4.5. Bayesian analysis

#### 2.4.6. Bayesian analyses were implemented in JASP

(JASP Team, 2019; Version 0.11.1) to estimate the likelihood ratio (Bayes factor) between the “no change in preference” and “change in preference” models

#### 2.4.7. Difference in WTP

Similar to the NHST, Bayesian two way repeated measures ANOVA was fitted to the data using a default prior r-scale of 0.5 (Rouder et al., 2012).

#### 2.4.8. Binary choice probe

Bayesian One Sample T-Test was fitted to the data using a default prior Cauchy scale of 0.707 (Rouder et al., 2009).

### 2.5. Data sharing

The datasets generated and analysed during the current study are available in the Open Science Framework (OSF) repository, https://osf.io/xfkgq/.

## 3. Results

### 3.1. perceived and required effort

There was a significant correlation between perceived and actual required effort in all four experiments (experiment 1, b = 0.95, t_(79)_ = 21.77, p<0.0001; experiment 2, b = 0.78, t_(103)_ = 19.51, p<0.0001; experiment 3, b = 0.85, t_(47)_ = 13.36, p<0.0001; experiment 4, b = 0.27, t_(47)_ = 5.14, p<0.0001).

### 3.2. Difference in WTP

#### 3.2.1. Null hypothesis significance testing

There was no difference in the mean Δbid between the different effort levels in any of the four experiments. There was a difference in the mean Δbid between the different value levels in all of the four experiments. High value items had negative mean Δbid (had lower value at the end of the experiment compared to the beginning) and low value items had positive mean Δbid (had higher value at the end of the experiment compared to the beginning). there was no interaction between effort and value levels in any of the four experiments.

#### 3.2.2. Bayesian analysis

In all 4 experiments comparing the effect of effort level in “no difference in WTP” and “difference in WTP” models resulted in large Bayes factor supporting the null (experiment 1, BF_null(effort)_= 1.89·10^8^; experiment 2, BF_null(effort)_= 4.74·10^8^; experiment 3, BF_null(effort)_= 2.77·10^5^; experiment 4, BF_null(effort)_= 1.43·10^12^).

### 3.3. Binary choice probe

#### 3.3.1. Null hypothesis significance testing

In both experiment 3 and 4, the choice ratio of higher effort over lower effort items was no different from chance (experiment 3, OR = 0.99 [0.86 1.14], z = −0.15, p = 0.88; experiment 4, OR = 0.96 [0.80 1.14], z = −0.48, p = 0.63; Figure 3).

#### 3.3.2. Bayesian analysis

In both experiment 3 and 4, comparing the models for change in choice ratio of higher effort items resulted in larger Bayes factor supporting the null (experiment 3, BF_null(choice)_= 4.23, median difference = −0.08 CI[-0.46 0.29]; experiment 4, BF_null(effort)_= 4.61, median difference = −0.02 CI[-0.40 0.34]).

## 4. Discussion

The aim of the current study was to examine whether effort association affects preferences. Simultaneous association of effort and appetitive items was used to probe preference change in humans. In all 4 experiments the participants were able to distinguish between the different effort levels and perform the effort association task. Across all 4 experiments we did not find a change in WTP or binary choice preference for similar value items following association of different levels of hand grip effort (Figures 2&3). In a Bayesian analysis we quantified the likelihood ratio of the null hypothesis (no change in preference) compared to alternative hypothesis (change in preference). This yielded very large Bayes factors in favor of the null (Table 4; Jeffreys, 1998).

**Figure 2.**
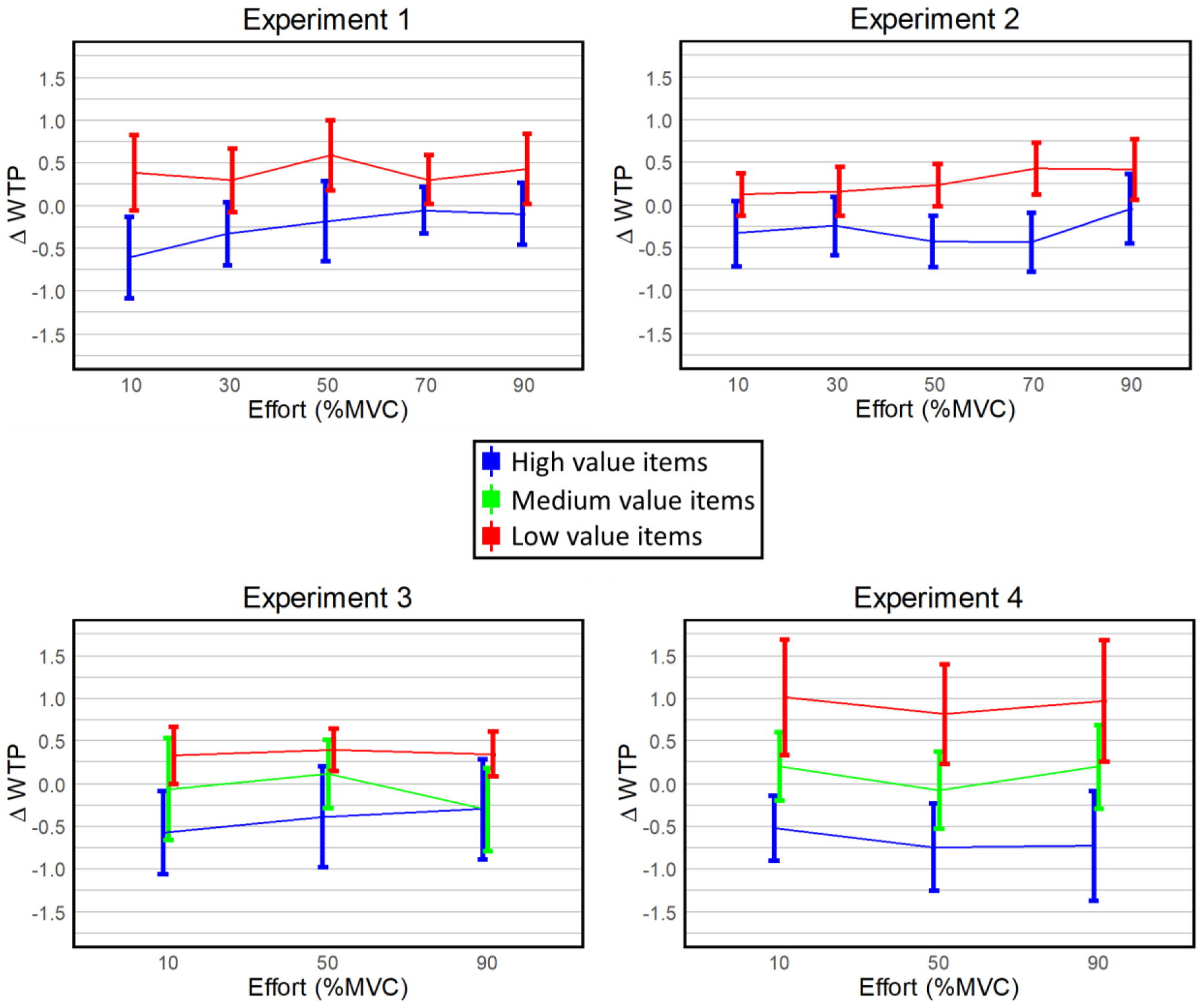
Change in WTP following effort association. Mean and CI of the difference in WTP (Δbid) for each value-effort category before and after effort association.

**Figure 3.**
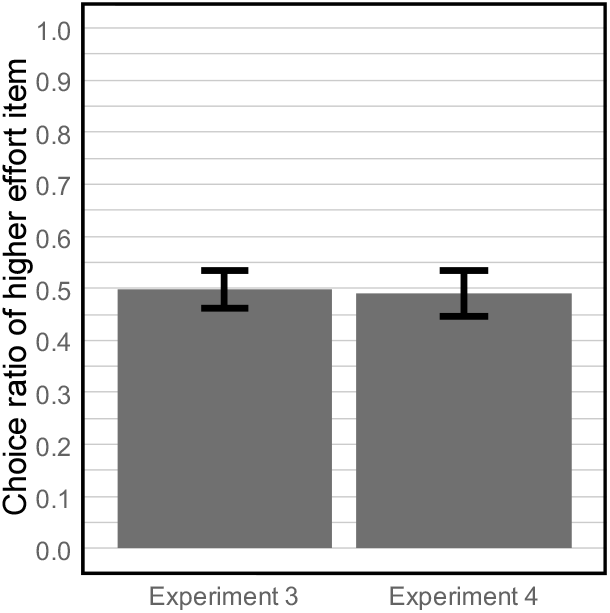
Probe. Choice ratio of similar initial value, higher effort over lower effort items.

**Table 4.** Bayesian analysis of effort and value on WTP

| Experiment | Model Comparison |  |  |  |  |
| --- | --- | --- | --- | --- | --- |
| | Models | P(M\|data) | $BF_{\text{Model}}$ | $BF_{\text{null}}$ | Error (%) |
| 1 | value | 0.87 | 27.32 | 1 |  |
| | effort | $4.63 \cdot 10^{-9}$ | $1.85 \cdot 10^{-8}$ | $1.89 \cdot 10^8$ | 1.53 |
|  | value*effort | 0.02 | 0.08 | 42.58 | 1.76 |
| 2 | value | 0.88 | 28.95 | 1 |  |
| | effort | $1.85 \cdot 10^{-9}$ | $7.41 \cdot 10^{-9}$ | $4.74 \cdot 10^8$ | 15.39 |
|  | value*effort | 0.01 | 0.05 | 69.74 | 2.88 |
| 3 | value | 0.92 | 44.42 | 1 |  |
| | effort | $3.35 \cdot 10^{-6}$ | $1.35 \cdot 10^{-5}$ | $2.77 \cdot 10^5$ | 1.28 |
|  | value*effort | 0.01 | 0.03 | 129.90 | 3.63 |
| 4 | value | 0.91 | 42.02 | 1 |  |
| | effort | $6.36 \cdot 10^{-13}$ | $2.54 \cdot 10^{-12}$ | $1.43 \cdot 10^{12}$ | 1.69 |
|  | value*effort | 0.003 | 0.01 | 312.73 | 2.34 |

Effort is typically aversive and weighted as a cost within decision making (Kurzban, 2016; Walton et al., 2006). Therefore, one might expect a negative effect of effort on the associated stimuli. Indeed a study with monkeys reported a preference for neutral stimuli that followed low effort over high effort (Shibasaki & Kawai, 2011). The stimuli in our study were appetitive snack food items. Therefore, the lack of a negative effect of effort could be due to the contradicting valence of the effort and the snack stimuli (Dickinson & Pearce, 1977). There are however indications that effort investment can lead to increase in preference toward the higher effort outcomes (for review see: Inzlicht et al., 2018). One of the most prevalent demonstrations for the effect of effort on preference is the within-trial contrast (WTC). In this paradigm increase in preference for item that follow effort expenditure is interpreted as resulting from the contrast between the negative affective state of effort and the relief of its termination (Zentall, 2005). The WTC effect has been repeatedly reported in birds, rodents and humans (Alessandri et al., 2008; Clement et al., 2000; Johnson & Gallagher, 2011; Kacelnik & Marsh, 2002; Lydall et al., 2010; Tsukamoto et al., 2017). However, studies have reported failures to replicate it (Arantes & Grace, 2008; Armus, 2001; Aw et al., 2011; Jellison, 2003; Shibasaki & Kawai, 2008; Vasconcelos et al., 2007; Vasconcelos & Urcuioli, 2008), and a study with monkeys reported an opposite effect (Shibasaki & Kawai, 2011). It has been suggested that insufficient training, prior history with lean schedules of reinforcement, and low statistical power may have been responsible for those failures (Zentall, 2008). Thus, further research is required to elucidate the robustness of this effect and the specific conditions that are required for effort to increase the value of associated items.

A major difference between WTC and our study is that instead of following effort production the stimuli in our study were presented simultaneously with effort production. This could explain the lack of increase in preferences toward higher effort items in our study. Furthermore, it supports the interpretation of WTC resulting from the contrast of relief from negative affective state of effort (Zentall, 2010). In contrast, the effort justification hypothesis suggests that outcomes are evaluated retrospectively according to the associated effort (Festinger & Carlsmith, 1959). According to this hypothesis, simultaneous presentation of effort and stimuli should not differ from sequential presentation. Although our study was not designed to replicate the WTC, it serves as evidence that mere association between effort demand and stimulus is not sufficient to induce preference change. Future studies could test whether the WTC effect result from the association of the relief from effort and the stimuli (backward conditioning) by presenting the stimulus prior to effort termination such that it predicts the relief from effort (forward conditioning) which should lead to an even larger effect (Escobar et al., 2004).

## Acknowledgements

This work was supported by the Israeli Science Foundation (ISF number 1798/15 and 2004/15) granted to Tom Schonberg.

## References

Alessandri, J., Darcheville, J.-C., Delevoye-Turrell, Y., & Zentall, T. R. (2008). Preference for rewards that follow greater effort and greater delay. Learning & Behavior : A Psychonomic Society Publication, 36(4), 352–358. https://doi.org/10.3758/LB.36.4.352

Arantes, J., & Grace, R. C. (2008). Failure to obtain value enhancement by within-trial contrast in simultaneous and successive discriminations. Learning and Behavior. https://doi.org/10.3758/LB.36.1.1

Armus, H. L. (2001). Effect of response effort on the reward value of distinctively flavored food pellets. Psychological Reports. https://doi.org/10.2466/pr0.88.4.1031-1034

Aw, J. M., Vasconcelos, M., & Kacelnik, A. (2011). How costs affect preferences: Experiments on state dependence, hedonic state and within-trial contrast in starlings. Animal Behaviour. https://doi.org/10.1016/j.anbehav.2011.02.015

Becker, G. M., Degroot, M. H., & Marschak, J. (1964). Measuring utility by a single???response sequential method. Behavioral Science, 9(3), 226–232. https://doi.org/10.1002/bs.3830090304

Bonnelle, V., Veromann, K. R., Burnett Heyes, S., Lo Sterzo, E., Manohar, S., & Husain, M. (2015). Characterization of reward and effort mechanisms in apathy. Journal of Physiology Paris. https://doi.org/10.1016/j.jphysparis.2014.04.002

Clement, T. S., Feltus, J. R., Kaiser, D. H., & Zentall, T. R. (2000). “work ethic” in pigeons: Reward value is directly related to the effort or time required to obtain the reward. Psychonomic Bulletin & Review, 7(1), 100–106. https://doi.org/10.3758/BF03210727

Colley, W. (2002). Colley’s bias free college football ranking method: The Colley matrix explained. Working Paper, 1–23.

Denk, F., Walton, M. E., Jennings, K. A., Sharp, T., Rushworth, M. F. S., & Bannerman, D. M. (2005). Differential involvement of serotonin and dopamine systems in cost-benefit decisions about delay or effort. Psychopharmacology, 179(3), 587–596. https://doi.org/10.1007/s00213-004-2059-4

Dickinson, A., & Pearce, J. M. (1977). Inhibitory interactions between appetitive and aversive stimuli. Psychological Bulletin. https://doi.org/10.1037/0033-2909.84.4.690

Escobar, M., Miller, R. R., Amundson, J., Arcediano, F., Chang, R., Frieda, E., Grace, R., Johnson, M., Parker, S., Stout, S., & Wheeler, D. (2004). A Review of the Empirical Laws of Basic Learning in Pavlovian Conditioning. In International Journal of Comparative Psychology.

Festinger, L. (1957). A theory of cognitive dissonance. In Scientific American (Vol. 207, p. 291). https://doi.org/10.1037/10318-001

Festinger, L., & Carlsmith, J. M. (1959). Cognitive consequences of forced compliance. Journal of Abnormal and Social Psychology. https://doi.org/10.1037/h0041593

Floresco, S. B., Tse, M. T. L., & Ghods-Sharifi, S. (2008). Dopaminergic and glutamatergic regulation of effort- and delay-based decision making. Neuropsychopharmacology : Official Publication of the American College of Neuropsychopharmacology, 33(8), 1966–1979. https://doi.org/10.1038/sj.npp.1301565

Hartmann, M. N., Hager, O. M., Tobler, P. N., & Kaiser, S. (2013). Parabolic discounting of monetary rewards by physical effort. Behavioural Processes. https://doi.org/10.1016/j.beproc.2013.09.014

Hull, C. L. (1943). Principles of Behavior: An Introduction to Behavior Theory. The Journal of Abnormal and Social Psychology, 39(3), 377–380. https://doi.org/10.1037/h0051597

Inzlicht, M., Shenhav, A., & Olivola, C. Y. (2018). The Effort Paradox: Effort Is Both Costly and Valued. In Trends in Cognitive Sciences. https://doi.org/10.1016/j.tics.2018.01.007

JASP Team. (2019). JASP. In [Computer software].

Jeffreys, H. (1998). The theory of probability. OUP Oxford.

Jellison, J. L. (2003). “Justification of effort” in rats: Effects of physical and discriminative effort on reward value. Psychological Reports. https://doi.org/10.2466/pr0.93.8.1095-1100

Johnson, A. W., & Gallagher, M. (2011). Greater effort boosts the affective taste properties of food. Proceedings. Biological Sciences / The Royal Society, 278(1711), 1450–1456. https://doi.org/10.1098/rspb.2010.1581

Kacelnik, A., & Marsh, B. (2002). Cost can increase preference in starlings. Animal Behaviour, 63(2), 245–250. https://doi.org/10.1006/anbe.2001.1900

Kurniawan, I. T., Seymour, B., Talmi, D., Yoshida, W., Chater, N., & Dolan, R. J. (2010). Choosing to make an effort: the role of striatum in signaling physical effort of a chosen action. Journal of Neurophysiology, 104(1), 313–321. https://doi.org/10.1152/jn.00027.2010

Kurzban, R. (2016). The sense of effort. Current Opinion in Psychology, 7, 67–70. https://doi.org/10.1016/j.copsyc.2015.08.003

Lewis, M. (1965). Psychological effect of effort. Psychological Bulletin, 64(3), 183–190. https://doi.org/10.1037/h0022224

Lydall, E. S., Gilmour, G., & Dwyer, D. M. (2010). Rats place greater value on rewards produced by high effort: An animal analogue of the “effort justification” effect. In Journal of Experimental Social Psychology (Vol. 46, Issue 6). https://doi.org/10.1016/j.jesp.2010.05.011

Phillips, P. E. M., Walton, M. E., & Jhou, T. C. (2007). Calculating utility: Preclinical evidence for cost-benefit analysis by mesolimbic dopamine. Psychopharmacology, 191(3), 483–495. https://doi.org/10.1007/s00213-006-0626-6

Plassmann, H., O’Doherty, J., & Rangel, A. (2007). Orbitofrontal cortex encodes willingness to pay in everyday economic transactions. The Journal of Neuroscience : The Official Journal of the Society for Neuroscience, 27(37), 9984–9988. https://doi.org/10.1523/JNEUROSCI.2131-07.2007

Prévost, C., Pessiglione, M., Météreau, E., Cléry-Melin, M.-L., & Dreher, J.-C. (2010). Separate valuation subsystems for delay and effort decision costs. The Journal of Neuroscience : The Official Journal of the Society for Neuroscience, 30(42), 14080–14090. https://doi.org/10.1523/JNEUROSCI.2752-10.2010

Rouder, J. N., Morey, R. D., Speckman, P. L., & Province, J. M. (2012). Default Bayes factors for ANOVA designs. Journal of Mathematical Psychology. https://doi.org/10.1016/j.jmp.2012.08.001

Rouder, J. N., Speckman, P. L., Sun, D., Morey, R. D., & Iverson, G. (2009). Bayesian t tests for accepting and rejecting the null hypothesis. In Psychonomic Bulletin and Review. https://doi.org/10.3758/PBR.16.2.225

Salamone, J. D. (1994). The involvement of nucleus accumbens dopamine in appetitive and aversive motivation. Behavioural Brain Research, 61(2), 117–133. https://doi.org/10.1016/0166-4328(94)90153-8

Shibasaki, M., & Kawai, N. (2008). The effects of response cost and time on choosing a stimulus. Shinrigaku Kenkyu. https://doi.org/10.4992/jjpsy.79.241

Shibasaki, M., & Kawai, N. (2011). The reversed work-ethic effect: Monkeys avoid stimuli associated with high-effort. Japanese Psychological Research. https://doi.org/10.1111/j.1468-5884.2010.00449.x

Skvortsova, V., Palminteri, S., & Pessiglione, M. (2014). Learning To Minimize Efforts versus Maximizing Rewards: Computational Principles and Neural Correlates. Journal of Neuroscience. https://doi.org/10.1523/JNEUROSCI.1350-14.2014

Stevens, J. R., Rosati, A. G., Ross, K. R., & Hauser, M. D. (2005). Will travel for food: Spatial discounting in two New World monkeys. Current Biology, 15(20), 1855–1860. https://doi.org/10.1016/j.cub.2005.09.016

Tsukamoto, M., Kohara, K., & Takeuchi, K. (2017). Effects of effort and difficulty on human preference for a stimulus: Investigation of the within-trial contrast. Learning and Behavior, 45(2), 135–146. https://doi.org/10.3758/s13420-016-0248-8

Vasconcelos, M., & Urcuioli, P. J. (2008). Deprivation level and choice in pigeons: A test of within-trial contrast. Learning and Behavior. https://doi.org/10.3758/LB.36.1.12

Vasconcelos, M., Urcuioli, P. J., & Lionello-DeNolf, K. M. (2007). FAILURE TO REPLICATE THE “WORK ETHIC” EFFECT IN PIGEONS. Journal of the Experimental Analysis of Behavior. https://doi.org/10.1901/jeab.2007.68-06

Von Neumann, J., & Morgenstern, O. (1944). Theory of Games and Economic Behavior. Princeton University Press, 625. https://doi.org/10.1177/1468795X06065810

Walton, M. E., Kennerley, S. W., Bannerman, D. M., Phillips, P. E. M., & Rushworth, M. F. S. (2006). Weighing up the benefits of work: Behavioral and neural analyses of effort-related decision making. Neural Networks, 19(8), 1302–1314. https://doi.org/10.1016/j.neunet.2006.03.005

Zentall, T. R. (2010). Justification of Effort by Humans and Pigeons: Cognitive Dissonance or Contrast? Current Directions in Psychological Science, 19(5), 296–300. https://doi.org/10.1177/0963721410383381

Zentall, Thomas R. (2008). Within-trial contrast: When you see it and when you don’t. In Learning and Behavior. https://doi.org/10.3758/LB.36.1.19

Zentall, Thomas R. (2005). A Within-trial Contrast Effect and its Implications for Several Social Psychological Phenomena. International Journal of Comparative Psychology, 18(4), 273–297.

